# Microvascular preservation and cardiomyocyte hyperplasia underlie adaptive right ventricular development in congenital heart disease-associated pulmonary arterial hypertension

**DOI:** 10.1101/2024.12.04.626551

**Authors:** Michael A. Smith, Eleana Stephanie Guardado, Jason Boehme, Sanjeev A. Datar, Emin Maltepe, Naveen Swami, Gary W. Raff, Aaron Bodansky, Joseph Moreno, Annelise Prince, Nevan Powers, Guo N. Huang, Vinicio de Jesus Perez, Jeffrey R. Fineman

## Abstract

**Background:** Right ventricular (RV) failure is the primary cause of death among patients with pulmonary arterial hypertension (PAH). Patients with congenital heart disease-associated PAH (CHD-PAH) demonstrate improved outcomes compared to patients with other forms of PAH, which is related to the maintenance of an adaptively hypertrophied RV. In an ovine model of CHD-PAH, we aimed to elucidate the cellular, microvascular, and transcriptional adaptations to congenital pressure overload that support RV function in CHD-PAH.

**Methods:** Fetal surgery was performed on late gestation lambs to insert a large aortopulmonary graft, leading to a persistent congenital left-right shunt and RV pressure load. At 3 days and 4-6 weeks of life, shunt RV microvasculature, cardiomyocyte structure, and myocardial growth mechanisms were compared to age-matched controls and unoperated fetal RV. RNA sequencing was performed to assess differences in the RV transcriptomes.

**Results:** At 4-6 weeks of age, shunt lambs demonstrate significant RV enlargement (shunt 37.1 ± 2.9g vs control 15.9 ± 1.0g, p<0.001) but maintain stable microvascular density (fetal 3.0 ± 0.6 vs shunt 2.9 ± 0.3 vs control 3.1 ± 0.6 capillaries per 1000 µm^3^, p>0.05). Shunt RV cardiomyocytes are significantly smaller by cross-sectional area and more numerous than age-matched controls (shunt 73.3 ± 5.5 µm^2^ vs control 99.2 ± 4.9 µm^2^, p=0.013). At 3 days, shunt RV cardiomyocytes show evidence of increased proliferative capacity and ongoing hyperplasia compared to controls. RNA sequencing analyses reveal a distinct gene expression profile in shunt RV consistent with a delay in terminal differentiation and metabolic adaptations to support adaptive function.

**Conclusions:** This study provides novel insights into the development of adaptive RV hypertrophy in CHD-PAH, demonstrating roles for preserved microvascular density and increased postnatal cardiomyocyte hyperplasia in supporting RV performance. Future investigations into the mechanisms underlying these changes could have significant implications for the development of novel therapeutic strategies for supporting RV function.

## Introduction

Right ventricular (RV) function is the strongest determinant of clinical outcomes in patients with pulmonary arterial hypertension (PAH).^1^ Patients with congenital heart disease-associated PAH (CHD-PAH) often demonstrate improved outcomes compared to patients with idiopathic or hereditary PAH, which appears to be related to an adaptive hypertrophy of the RV.^2–4^ This is best illustrated in patients who develop Eisenmenger’s Syndrome, who often generate increased RV afterload to a similar degree as patients with other forms of PAH but are able to maintain a concentrically hypertrophied RV with preserved function until very late stages of disease.^5–8^

Little is known about the mechanisms underlying the development of the adaptively hypertrophied RV seen in patients with CHD-PAH. In utero, the RV is the dominant ventricle, providing roughly 60% of the total cardiac output to the fetal circulation.^9,10^ An increasingly dense capillary bed feeds the fetal myocardium as it grows primarily via cellular hyperplasia until terminal differentiation of cardiomyocytes occurs in the late gestation and early postnatal period.^11–14^ After the transition to postnatal circulation, in normal cardiac development the RV remodels to become a thin-walled, compliant compartment and further postnatal growth of the myocardium occurs primarily via cellular hypertrophy.^15^ It is unclear how the abnormal forces introduced by congenital heart lesions that lead to the development of CHD-PAH and RV overload may alter the typical developmental remodeling undertaken by the RV.

In an ovine model of CHD-PAH created via fetal implantation of an aortopulmonary shunt, we have previously characterized the physiologic performance of the hypertrophied RV that develops in response to chronic left-to-right shunting and elevated pulmonary arterial pressures.^16,17^ The shunt RVs demonstrate a unique capacity to maintain cardiac output in the face of acute afterload via augmenting myocardial contractility in a manner not seen in control animals. In the present work, we sought to characterize the cellular, microvascular, and transcriptional changes associated with the development of this adaptively performing RV in CHD-PAH.

## Methods

### Lamb model of CHD-PAH with adaptive RV hypertrophy

The study design is summarized in Figure 1. Fetal surgery was performed on late gestation mixed-breed Western lambs to insert an 8.0mm GoreTex graft (shunt) anastomosing the ascending aorta and the main pulmonary artery. This procedure has been described in detail previously.^16,17^ In brief, pregnant ewes were anesthetized and fetal exposure was obtained through the uterine horn. A left lateral thoracotomy was performed on the fetal lamb, through which the shunt was placed with the use of side biting vascular clamps. Lambs were then returned to the uterus and allowed to deliver spontaneously at term gestation roughly 1 week later. Control lambs were either provided by twin gestation or age matched. Fetal lamb hearts were harvested after elective cesarean delivery at 135-140 days gestation (term lamb gestation: ∼148 days).

**Figure 1:**
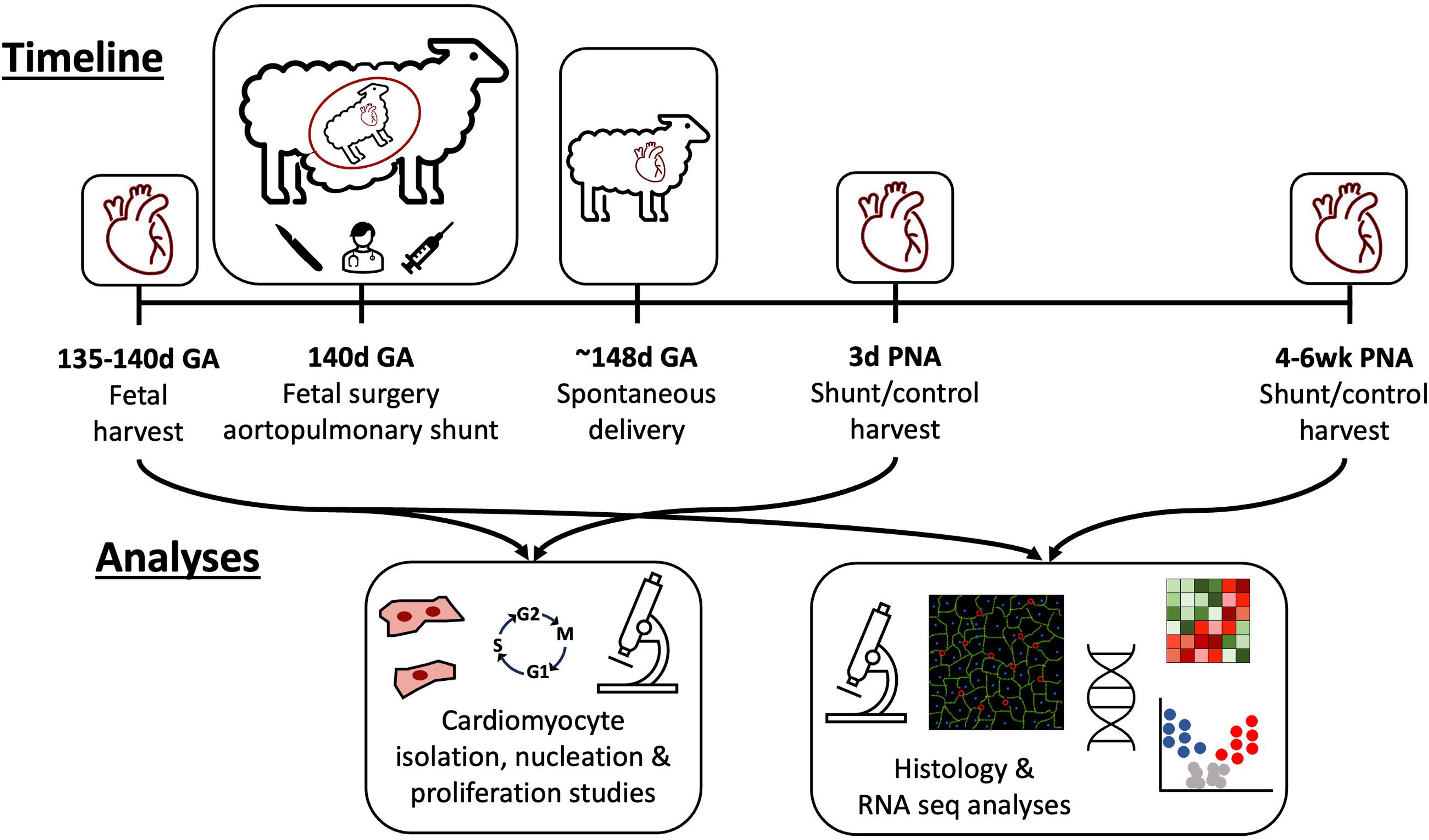
Schematic of study design Fetal surgery to implant an aortopulmonary graft was performed at 140 days gestational age (GA) in shunt lambs. Following spontaneous delivery, right ventricular tissue was harvested from shunt and control lambs at 3 days and 4-6 weeks postnatal age (PNA). Fetal tissue was obtained at 135-140 days GA. Cardiomyocyte isolation and nucleation/proliferation studies were performed on fetal and 3 day old shunt and control right ventricles. Myocardial structural histology and RNA sequencing analyses were performed on fetal and 4-6 week shunt and control right ventricles.

At three days and four to six weeks after delivery, lambs were anesthetized and instrumented for tissue harvest. Prior to harvest, lambs were euthanized with sodium pentobarbital followed by bilateral thoracotomy as described in the NIH Guidelines for the Care and Use of Laboratory Animals. The heart chambers were dissected into separate atria, RV free wall, and left ventricle plus septum (LV+S) segments which were then weighed. Full thickness sections of the RV free wall were then sampled and preserved or snap frozen. All protocols and procedures relating to the care and evaluation of these animals were approved by the Institutional Animal Care and Use Committees (IACUC) of the University of California, San Francisco and the University of California, Davis.

### Histologic assessment of RV free wall and isolated cardiomyocytes

RV free wall was fixed in 4% paraformaldehyde and embedded in optimal cutting temperature compound, then sectioned at 5-10µm for immunofluorescent staining. Sections were incubated with a permeabilizing blocking buffer for 1 hour followed by diluted primary antibody solutions overnight at 4 °C and secondary antibody incubation for 90 minutes (see Supplemental Table 1 for antibody, reagent, and buffer details). Slides were then mounted with 4’,6-diamidino-2-phenylindole (DAPI)-containing mounting medium to stain nuclei. Isolated cardiomyocytes were obtained as previously described from tissue segments that were fixed in 4% paraformaldehyde then incubated overnight in 50% w/v potassium hydroxide solution.^18^ The RV tissue was then briefly washed with PBS and gently crushed to release dissociated cardiomyocytes. Cells were then further washed 3 times with PBS prior to incubation with primary antibody solutions overnight. After a 90 minute secondary antibody incubation, cells in solution were deposited onto slides, allowed to dry completely, and mounted with DAPI-containing mounting medium. Images of stained sections and isolated cardiomyocytes were obtained using an Olympus IX51 fluorescent microscope equipped with a CCD camera (Hamamatsu Photonics) and processed and analyzed in Fiji/ImageJ and CellProfiler.^19–21^

### RNA sequencing and transcriptome analyses

RNA sequencing was performed on snap frozen tissue obtained from the RV free wall of fetal and 4-6 week old control and shunt lambs. RNA was isolated using the RNeasy Plus Universal Mini (QIAGEN) according to manufacturer’s instructions. Library preparation was done using the QIAseq Stranded mRNA Kit (QIAGEN). After quality control, sequencing was performed on a NextSeq instrument (2×75bp 2×10) according to the manufacturer instructions (Illumina Inc.) Reads were trimmed and ribosomal RNA reads were removed. All remaining reads were then mapped to the Sheep (Texel) genome Ovis Aries (Ovar v3.1) with ENSEMBL version 103 annotation. On average, 66 million paired-end reads per sample were mapped with an average mapping rate of 77%.

Unsupervised cluster and differential gene expression analyses were performed using the EdgeR pipeline in R, version 4.2.1.^22,23^ Differentially expressed genes with false discovery rate (FDR) <0.01 based on Benjamini Hochberg algorithm and absolute fold change ≥1.2 were considered statistically significant. Gene Ontology (GO) enrichment analyses were performed using the *ClusterProfiler* package.^24^ Human transcriptome data was derived from previously published work and downloaded from Gene Expression Omnibus (GSE240941).^25^ Gene set variation analysis was performed with the *GSVA* package using GO sheep genome gene sets derived from AnnotationHub record AH107722.^26–28^

### Statistical analyses

Data are presented as mean ± standard error. Baseline lamb characteristics were compared between shunt and control animals using unpaired T tests for continuous data and Fisher’s exact test for categorical data. Microvascularity metrics, cardiomyocyte area and density, and cardiomyocyte nucleation and proliferation rates were compared using linear mixed effects models incorporating an animal identifier as a random effect to account for the correlation structure of technical replicates.^29,30^ Enrichment scores derived from gene set variation analysis were compared using Kruskal Wallis tests for multi-group comparisons. T tests were used for targeted pairwise comparisons. A p value <0.05 was considered statistically significant. All statistical analyses were performed in R, version 4.2.1.^23^

## Results

### Animal Details

Hearts from late gestation fetal, 4-6 week old shunt, and 4-6 week old control lambs were evaluated for the primary analyses of microvascular and cardiomyocyte architecture and transcriptomics. Details of lamb age, sex, and heart chamber weights for the 4-6 week old shunt and control lambs are provided in Table 1. The shunt lambs were marginally younger than control at time of harvest (30.9 ± 0.9 days vs 35.6 ± 1.2, p=0.007). Shunt lambs had lower body weight but enlarged hearts compared to control, including by total heart weight, weight of left ventricle plus septum, and weight of right ventricle. These significant differences persisted with adjustment for body weight.

**Table 1:**
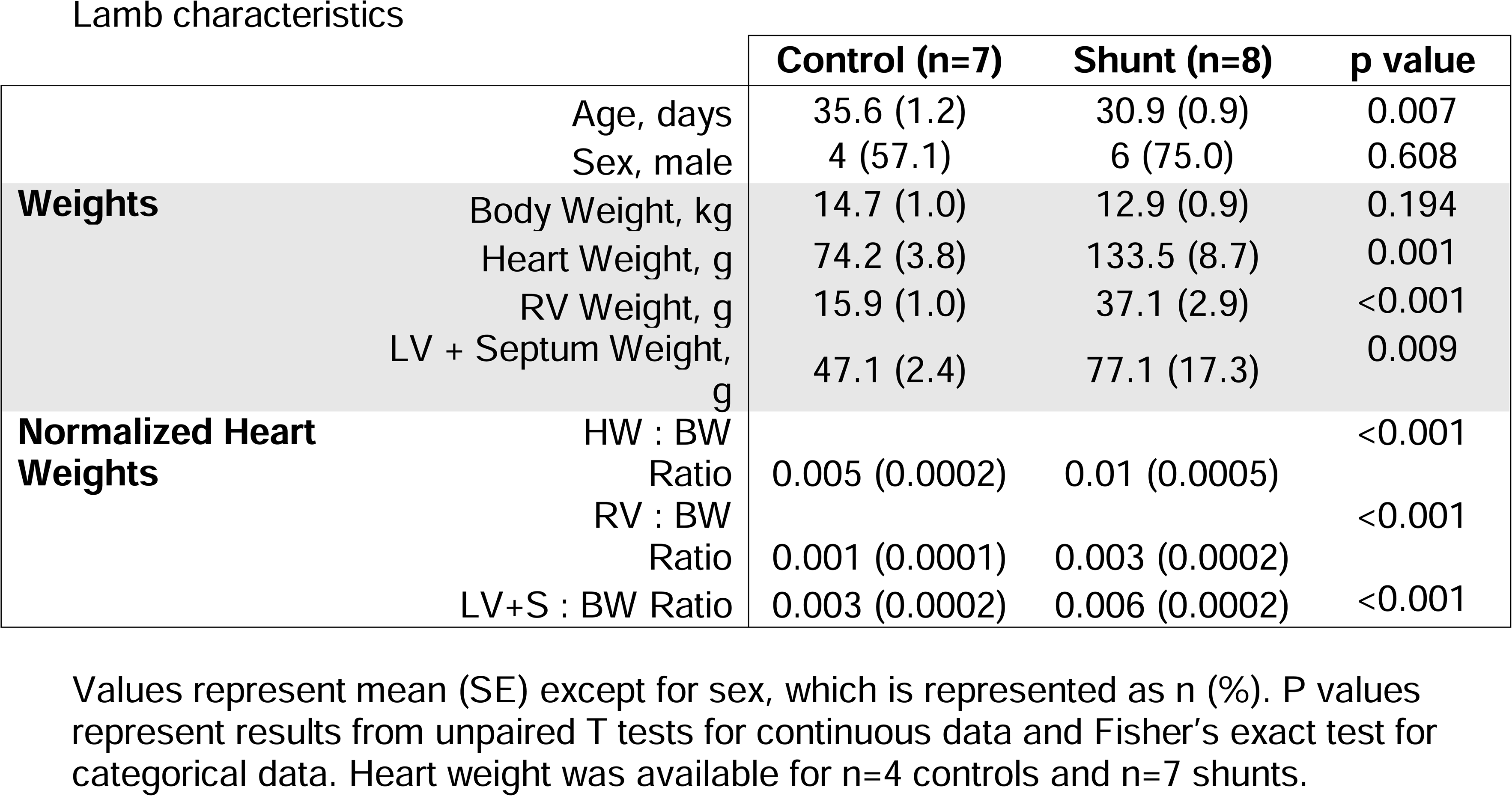
Lamb characteristics.

### Microvascular Architecture

The microvascular architecture of the RV was examined through histologic assessment of the myocardial capillary bed. The microvasculature was quantified using two metrics, capillary density and capillary per cardiomyocyte ratio (Figure 2). The average capillary density did not significantly differ between late gestation fetal, 4-6 week old shunt, and 4-6 week old control RV myocardium (3.0 ± 0.6, 2.9 ± 0.3, and 3.1

**Figure 2:**
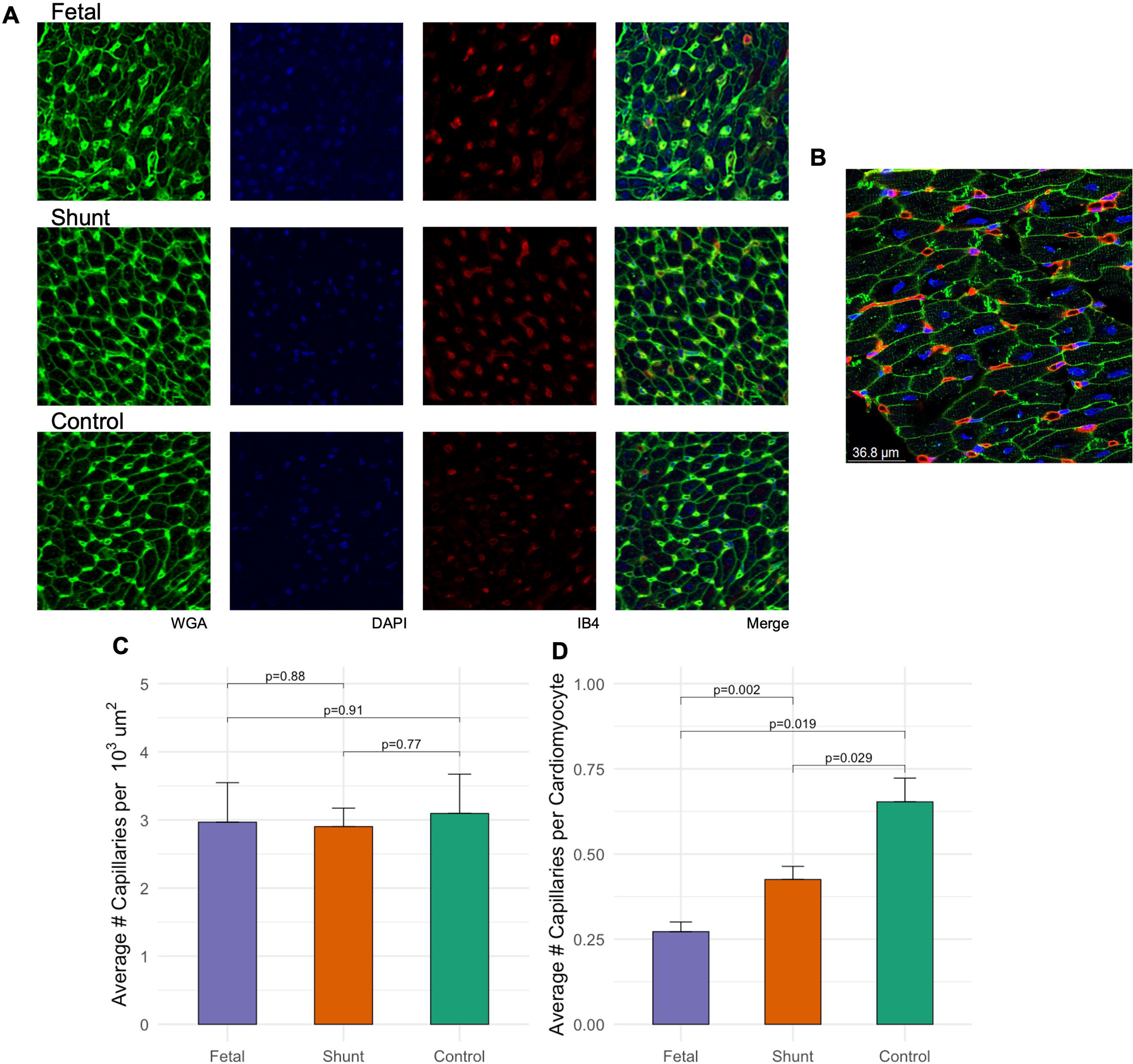
Right ventricular microvascular architecture A) Right ventricle myocardium from fetal and 4-6 week old shunt and control lambs was stained with wheat germ agglutinin (WGA, green) to delineate cardiomyocyte cell walls, 4′,6-diamidino-2-phenylindole (DAPI, blue) for nuclei, and isolectin B4 (IB4, red) to delineate endothelial cells. B) 60x magnification demonstrates the capillary and cardiomyocyte structure in detail. C) Capillary density and D) capillary to cardiomyocyte ratios were compared between fetal and 4-6 week old shunt and control right ventricles.

± 0.6 capillaries per 1000 µm^3^ respectively, p>0.05, Figure 2C). Capillary per cardiomyocyte ratios were significantly different between the 3 groups, however. The average number of capillaries per cardiomyocyte in fetal tissue (0.27 ± 0.03) was significantly lower than in shunt myocardium (0.43 ± 0.04, p=0.002) and in control myocardium (0. 65 ± 0.07, p=0.019). In comparisons between shunt and control myocardium, the capillary per cardiomyocyte ratio was significantly smaller in shunt RV (p=0.029, Figure 2D). The differences in capillary per cardiomyocyte ratios in the setting of stable capillary densities suggest differences in the size and density of cardiomyocytes. Thus, the cardiomyocyte structure was assessed next.

### Cardiomyocyte Structure

Average cardiomyocyte transverse cross-sectional areas and density of cardiomyocytes were compared (Figure 3). Cardiomyocytes were significantly smaller by cross-sectional area in the fetal RV (30.6 ± 3.9 µm^2^) compared to 4-6 week shunt (73.3 ± 5.5 µm^2^, p<0.001) and control RV (99.2 ± 4.9 µm^2^, p<0.001). The shunt RV cardiomyocytes were significantly smaller than age-matched control (p=0.013, Figure 3B). The expected inverse relationships were noted in cardiomyocyte density, with the fetal RV having the highest density of cardiomyocytes, followed by shunt and finally control myocardium (p<0.05 for all comparisons, Figure 3C). Compared to age-matched controls, the enlarged shunt RV myocardium appears to consist of more numerous, smaller cardiomyocytes.

**Figure 3:**
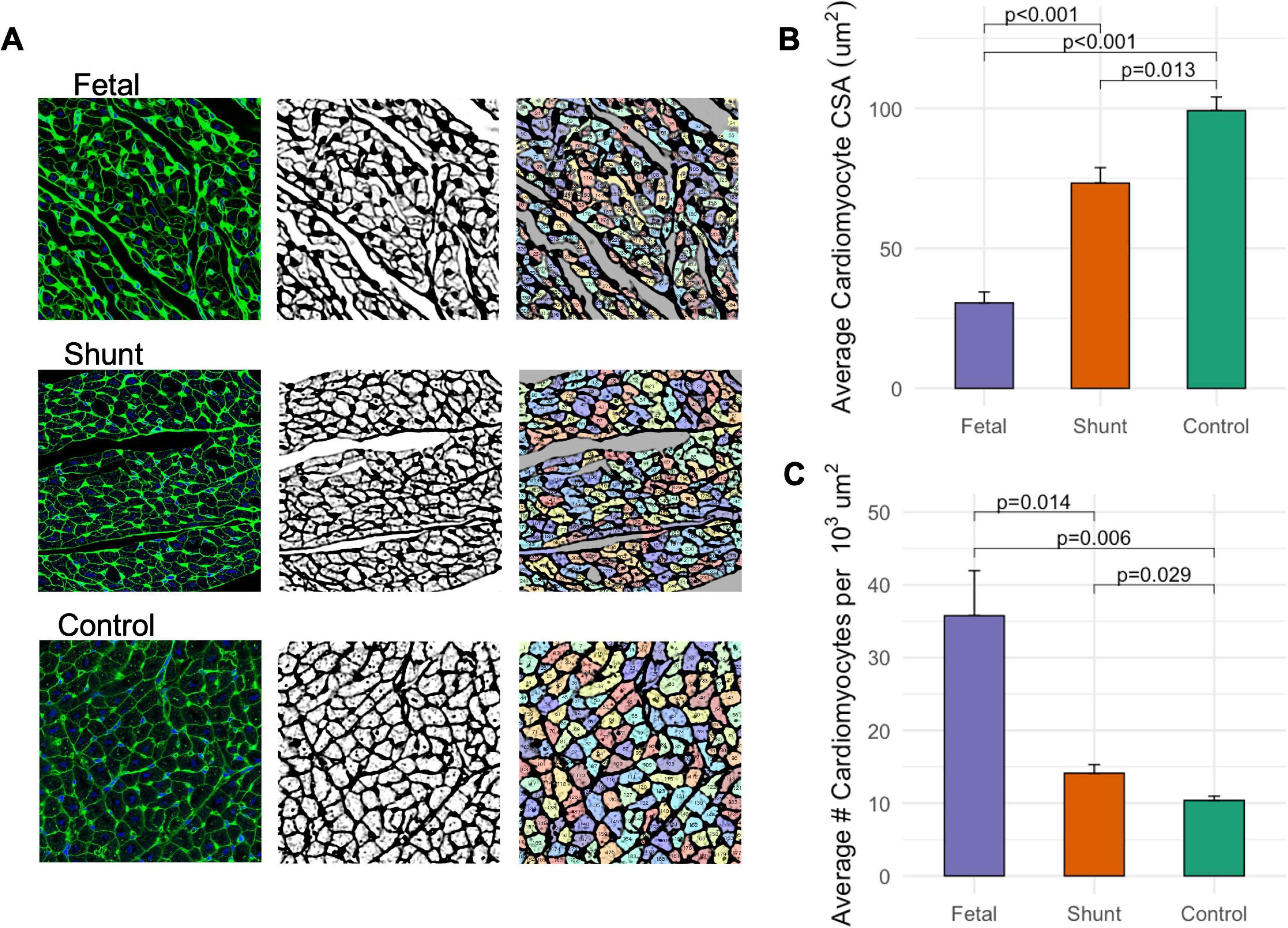
Right ventricular cardiomyocyte structure A) Wheat germ agglutinin-stained images of the cardiomyocyte cell walls were processed and segmented to determine the transverse cross sectional areas (CSA) of cardiomyocytes. B) Cardiomyocyte CSA and C) density of cardiomyocytes were compared between fetal and 4-6 week old shunt and control right ventricles.

### RV Transcriptomic Signatures

To investigate the transcriptional differences pertinent to the unique myocardial architecture of the shunt RV, we performed bulk RNA sequencing of RV free wall tissue from fetal and 4-6 week shunt and control RV. In principal components analysis and unsupervised hierarchical clustering of gene expression data, we found that fetal, shunt, and control RVs cluster by distinct transcriptomes (Figure 4). Differential gene expression analyses were performed to highlight the distinct developmental trajectories undertaken by shunt and control RV. Compared to fetal RV, there were 560 genes significantly upregulated and 415 genes downregulated in shunt RV and 226 genes upregulated and 253 downregulated in control RV (FDR < 0.01 and absolute fold change ≥ 1.2, Figure 4C, Supplemental Table 2, Supplemental Figure 1). There were 427 and 276 uniquely up- and downregulated genes, respectively, during development in shunt RV versus just 93 up- and 114 downregulated genes in controls, highlighting the unique developmental trajectory undertaken by shunt RV (Figure 4D). Comparing shunt to control RV, there were 100 significantly upregulated and 96 downregulated genes.

**Figure 4:**
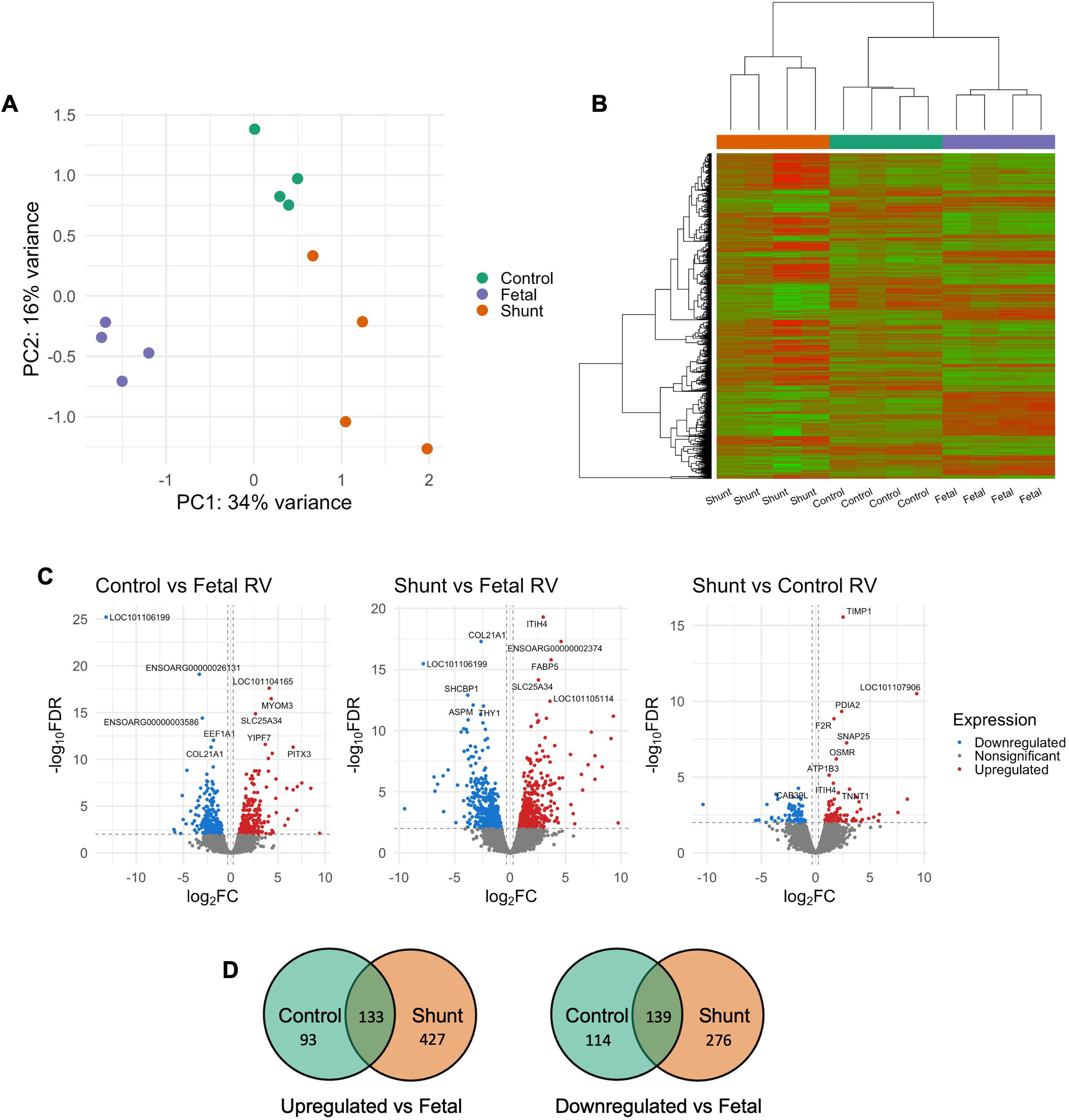
RNA sequencing of right ventricle myocardium Right ventricular tissue from fetal and 4-6 week old shunt and control lambs underwent RNA isolation and sequencing. A) Principal components analyses and B) hierarchical clustering of relative gene expression levels demonstrate distinct clustering of the fetal, shunt, and control right ventricle transcriptomes. C) Volcano plots depict results of pairwise differential gene expression analyses. D) Venn diagrams highlight the differences in developmental trajectories between shunt and control right ventricles, with notably more genes up- and down-regulated in shunts than controls vs fetal.

The genes differentially expressed between shunt and control RV are highlighted in Figure 5. In GO enrichment analyses, the genes differentially regulated in shunt RV enrich pathways primarily involved in mitochondrial energy metabolism, aerobic respiration, and the electron transport chain (Figure 5B). These results were compared to GO enrichment analyses of previously published human transcriptome data evaluating RV tissue from patients with compensated RV hypertrophy and patients with decompensated or failing RVs (Figure 5C).^25^ In this study, patients with compensated hypertrophy demonstrated preserved cardiac indices (>2.2 L/min/m^2^) and/or normal tricuspid annular plane systolic excursion (TAPSE) on echocardiography (≥17mm). Decompensated RV tissue was obtained from patients with PAH or dilated cardiomyopathy who received heart/lung transplant or died with RV failure. Several of the differentially enriched biological processes differentiating compensated and decompensated RVs in humans were similar to those observed differentiating shunt and control RV. Specifically, oxidative phosphorylation and the mitochondrial respiratory chain are important to supporting RV function in both compensated human RV and in our CHD-PAH shunt model.

**Figure 5:**
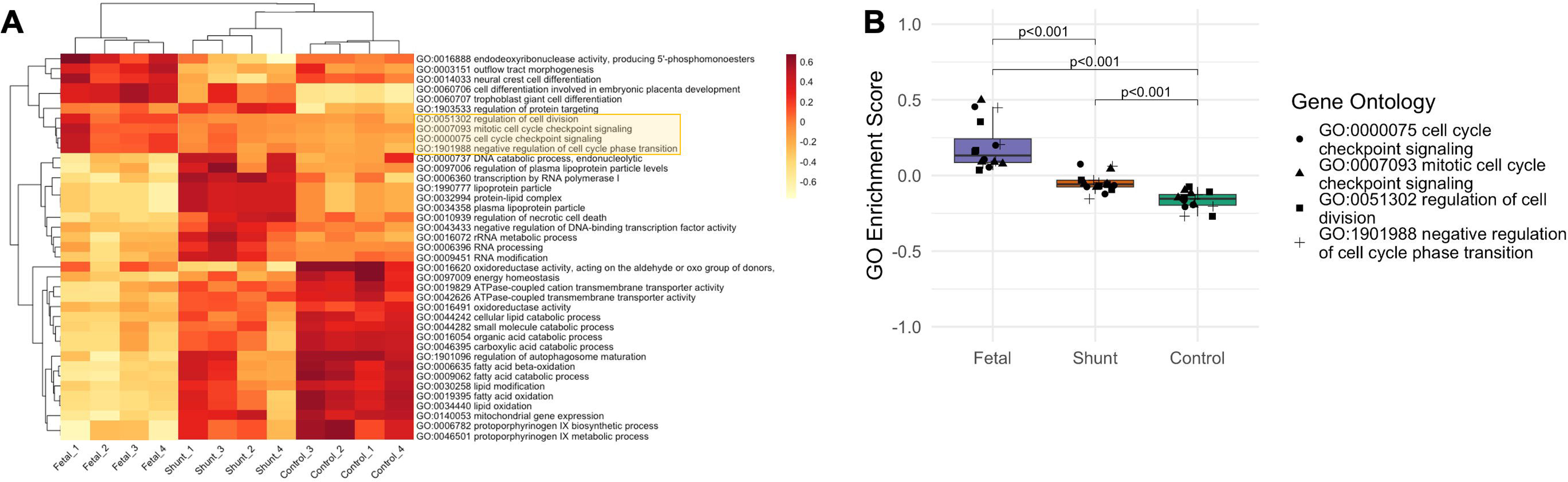
Shunt vs control right ventricle transcriptome and comparison to humans A) Heatmap of relative expression levels of the 196 genes significantly differentially expressed between shunt and control right ventricles. B) The genes differentially expressed between shunt and control enrich Gene Ontologies pertinent to mitochondrial energy metabolism, aerobic respiration, and the electron transport chain. Gene counts indicate the number of genes differentially expressed that are annotated to a given ontology, gene ratio indicates the gene count divided by the total number of differentially expressed genes, and p values are derived from Fisher’s exact tests adjusted for multiple comparisons. C) Gene Ontology enrichment analyses of previously published human right ventricle transcriptome data comparing compensated and decompensated ventricles demonstrate similarly enriched pathways (highlighted in yellow).

Gene set variation analysis was utilized to better understand the biological processes represented by the differential gene expression profiles of the fetal, shunt, and control RV. Figure 6A depicts the relative enrichment scores of GO biological processes that were significantly differentially enriched between the fetal, shunt, and control groups. Hierarchical cluster analysis of the biological process enrichment scores again distinctly clusters the 3 groups of animals. Four biological processes that were differentially enriched related to the cell cycle, specifically negative regulation of cell cycle progression. These processes were activated in the late gestation fetal myocardium, presumably driving the terminal differentiation of cardiomyocytes known to occur in this time period. These processes were negatively enriched, or suppressed, in the 4-6 week old control RV, which consists of mature differentiated cardiomyocytes. The shunt RV demonstrated an intermediate enrichment score, suggesting a potential delay in the terminal differentiation process (Figure 6B).

**Figure 6:**
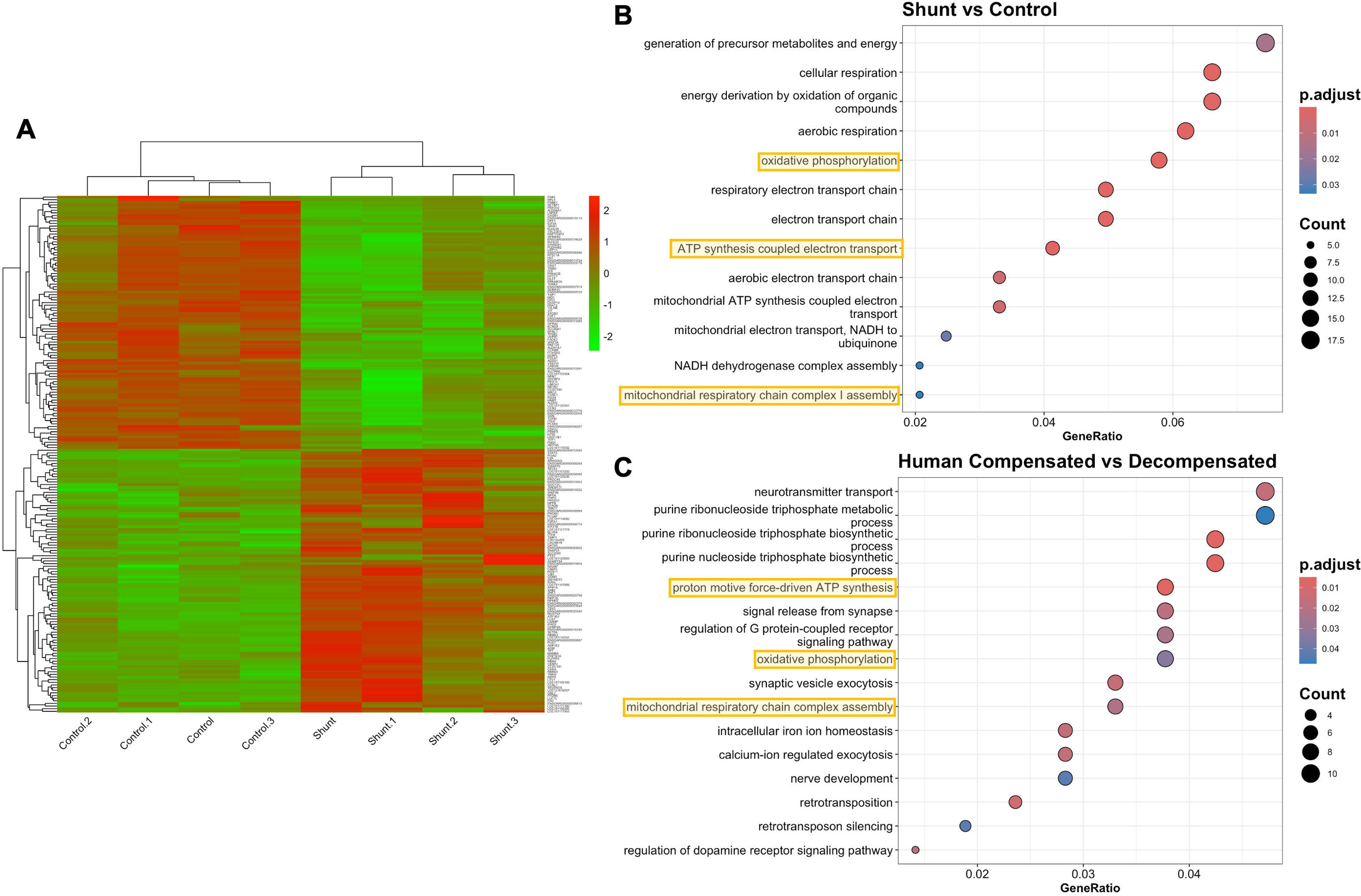
Gene set variation analyses of fetal, shunt, and control right ventricle transcriptomes A) Heat map of relative enrichment scores of the Gene Ontology biological processes that were significantly differentially expressed between fetal, shunt, and control right ventricle. B) Four pathways relating specifically to negative regulation of the cell cycle and terminal differentiation were differentially enriched between fetal, shunt, and control right ventricle.

### Cardiomyocyte Nucleation and Proliferation in Early Postnatal Life

We next further investigated the potential for increased cardiomyocyte proliferation or delayed terminal differentiation contributing to the development of an overall enlarged myocardium consisting of smaller, more densely packed cardiomyocytes in the shunt animals. Proliferative capacity of cardiomyocytes is known to be relatively restricted to the fetal and early postnatal periods.^31^ Therefore, myocardium from 3 day old shunt and control animals was examined for the following analyses. Terminal differentiation of cardiomyocytes often results in bi- or polynucleated mature cardiomyocytes, and thus mononucleate cells can be quantified as a surrogate for cells with proliferative capacity (Figure 7A and 7B).^12,13^ At 3 days of age, the percentage of mononucleate cardiomyocytes in shunt RV (32.9 ± 1.8%) was similar to that in fetal RV (34.9 ± 1.1%, p=0.85) and significantly greater than 3 day old control RV (15.2 ± 1.4%, p<0.001).

**Figure 7:**
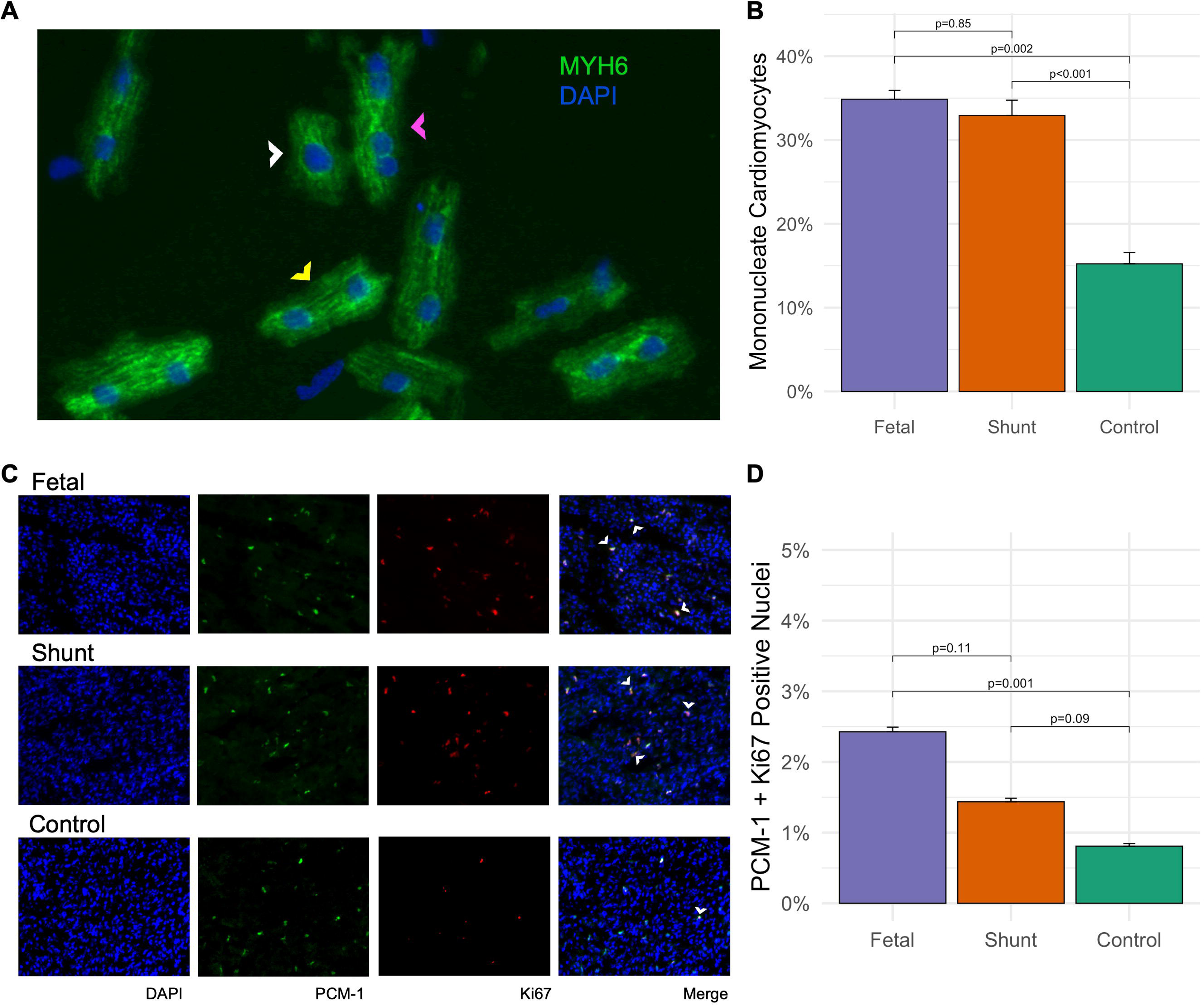
Cardiomyocyte nucleation and proliferation in early post-natal life Cardiomyocytes were isolated from fetal and 3 day old shunt and control right ventricle. A) Immunofluorescent staining of alpha-myosin (MYH6) and nuclei (DAPI) demonstrates the presence of mono- (white arrow), bi- (yellow arrow), and poly- nucleated (pink arrow) cardiomyocytes. B) The percentage of mononucleate cardiomyocytes, an surrogate for cells with proliferative capacity, was compared between fetal and 3 day old shunt and control right ventricle. C) Cardiomyocyte cell cycle activity was assessed through immunostaining of nuclei (DAPI), pericentriolar marker-1 (PCM-1), and Ki67. Cardiomyocytes active in the cell cycle are indicated with white arrows. D) The percentage of PCM-1 + Ki67 positive nuclei was compared between fetal and 3 day old shunt and control right ventricle.

To evaluate the proportion of cardiomyocytes active in the cell cycle at 3 days of life, RV tissue sections were stained with pericentriolar marker 1 (PCM-1), a mature striated muscle-specific perinuclear marker, and Ki67, a nuclear protein present only during the active phases of the cell cycle (Figure 7C and 7D). Fetal RV showed a significantly higher percentage of PCM-1 + Ki67 positive nuclei compared to 3 day old control RV (2.4 ± 0.07% vs 0.8 ± 0.04%, p<0.001). The percentage of PCM-1 + Ki67 positive nuclei in shunt RV (1.4 ± 0.05%) was intermediate between the fetal and control RV (p=0.11 vs fetal and p=0.09 vs control).

## Discussion

Utilizing a large animal model of CHD-PAH, we present novel insights into the microvascular and cardiomyocyte architecture of an adaptively performing, hypertrophied right ventricle. These results present a potential structural basis for the improved right ventricular function and clinical outcomes seen in patients with CHD-PAH and Eisenmenger’s Syndrome. Our shunt model imposes a persistent pressure load on the pulmonary vasculature and RV similar to what occurs with post-tricuspid congenital heart lesions such as a ventricular septal defect or patent ductus arteriosus. This prohibits the typical hemodynamic transition that occurs at birth and allows for what appears to be a maintenance of the fetal angiogenic and proliferative phenotype in the RV. The result is a hypertrophied RV with a preserved capillary density and increased number of cardiomyocytes, the contractile units of the ventricle. Together, these structural alterations appear to support the adaptive performance of the shunt RV.

Microvascular insufficiency is a key feature of maladaptive ventricular hypertrophy. In pathologic RV remodeling, impaired angiogenesis and capillary rarefaction lead to microvascular ischemia and subsequent myocardial injury that drives RV decompensation.^32^ Despite the significant hypertrophy of the myocardium in our shunt animals, capillary density was maintained at a level similar to the late gestation and age-matched control tissue as well as to previous reports in the literature,^33^ implying an angiogenic response to the postnatal pressure stimulus of the shunt. RV angiogenesis has similarly been described as an adaptive response to several other RV stressors, including chronic hypoxia-induced pulmonary hypertension and in a model of RV hypertrophy induced by pulmonary arterial banding.^34,35^ This is the first study to our knowledge to demonstrate such a response to the abnormal pressures and flow forces created by a congenital left-to-right shunt lesion. Interestingly, with the changes observed in cardiomyocyte size and density in our shunt RV, the capillary to cardiomyocyte ratio decreased, a feature sometimes seen in models of ventricular maladaptation.^36^ Reconciling this finding with the preserved capillary density and adaptive performance of these ventricles leads us to consider the efficiency of oxygen and substrate utilization at the cellular level. As evidenced by the results of GO enrichment analyses of our shunt vs control RV as well as in human compensated vs decompensated RV, maintaining mitochondrial energy metabolism and aerobic respiration appear critical to adaptive RV function. Further investigation is warranted to determine the metabolic efficiency and more thoroughly characterize the vascular supply of these adaptively performing cardiomyocytes.

With our findings of increased cardiomyocyte cellularity in the enlarged RV myocardium, we propose a novel structural mechanism that may contribute to the adaptive function of the RV seen in CHD-PAH and Eisenmenger Syndrome. Cardiomyocyte hyperplasia is the predominant growth mechanism in early gestation but it is gradually replaced by cellular hypertrophic growth as cells terminally differentiate around term gestation.^12–14^ Further proliferative capacity is severely limited in mature cardiomyocytes.^31,37^ Recently, several studies have demonstrated a hyperplastic response to several ventricular stimuli early in postnatal life, including direct traumatic injury and coronary artery ligation.^15,38,39^ Increased myocardial pressure loading of the ventricle, as is experienced in post-tricuspid left-right shunts, may act as a similar stimulus. Prior studies have demonstrated that increasing pressure on the developing ventricles via intrauterine aortic or pulmonary arterial banding led to increased cardiomyocyte proliferation.^40–42^ Postnatally, transverse aortic constriction triggered increased left ventricular cardiomyocyte proliferation in mice, though only when performed in the first few days after birth.^43^ The aortopulmonary shunt placed in our model exposes the right ventricle to a persistent systemic pressure load starting in that critical time period immediately following delivery. This appears to have led to a persistent proliferative phenotype in early postnatal life along with significant transcriptional changes and evidence of delayed terminal differentiation in our transcriptomic data as late as 4-6 weeks of age. We suspect that the resultant increased number of cardiomyocytes is a primary factor in the physiologic performance of these ventricles, as myocyte number has been shown to correlate with the capacity of the heart to adapt to increased hemodynamic loads.^12,44^

Further characterization of the molecular processes driving these findings could have significant clinical and therapeutic ramifications. First, while the majority of clinical studies demonstrate survival advantages for patients with CHD-PAH compared to other forms of PAH,^2–4^ results are not uniform across all investigations.^4,45^ There is significant heterogeneity amongst patients with CHD-PAH and refined phenotyping based on improved understanding of the physiologic mechanisms determining RV performance could help to improve classification and prognostication efforts. Additionally, our findings support the further investigation of models of CHD associated with abnormal pressure loads to understand the mechanisms underlying postnatal cardiomyocyte proliferation and the potential for myocardial regeneration. Better understanding of these mechanisms could unveil new therapeutic targets for supporting ventricular function or even regenerating injured myocardium in a wide array of diseases, including pulmonary hypertension, post-myocardial infarction, cardiomyopathies, and more.

There are several limitations to our work worth noting. First, our model best represents early-stage CHD-PAH and right ventricular remodeling but does not reflect more advanced disease at the time points studied. Evaluations of microvascular and cardiomyocyte architecture were limited to 2 dimensional assessments and further work to characterize the myocardium in 3 dimensions is merited. In our transcriptional profiling, we describe differential expression and enrichment analyses from bulk RNA sequencing. While most myocardial cells are cardiomyocytes, we cannot differentiate the expression profiles of all cell types in the myocardium from the present data. Nonetheless, the insight provided from these analyses provide ample evidence for meaningful structural and transcriptional changes related to shunt physiology.

In conclusion, in our ovine model of CHD-PAH we identified a unique developmental trajectory of the RV characterized by the maintenance of microvascular density and increased cardiomyocyte proliferative growth in addition to cellular hypertrophic growth. These structural adaptations could plausibly underlie the adaptive physiologic performance of the RV in CHD-PAH via preserving oxygen and substrate delivery and increasing the stock of contractile units within the myocardium. Further investigation to identify the specific mechanisms underlying these changes could have significant implications for supporting RV function not only in PAH but in any disease associated with ventricular failure.

---

Supplemental Figure 1: Normalized expression levels of top differentially expressed genes Normalized log_2_ counts per million (log_2_CPM) of the top 10 differentially expressed genes (by lowest false discovery rate) from pairwise comparisons of control vs fetal, shunt vs fetal, and shunt vs control analyses.

