## Supplemental Table 1 for "Microvascular preservation and cardiomyocyte hyperplasia underlie adaptive right ventricular development in congenital heart disease-associated pulmonary arterial hypertension"

Antibodies, Reagents, and Buffers

Wheat germ agglutinin Alexa Fluor 488 (1:300) (Thermo Fisher Scientific W11261)

Isolectin GS-IB4 Alexa Fluor 594 (1:150) (Thermo Fisher Scientific Invitrogen I21413)

VECTASHIELD PLUS Antifade Mounting Medium with DAPI (Vector Laboratories NC1848444)

Mouse anti-MYH6 antibody (1:500) (Abcam ab207926)

Rabbit anti-PCM1 (H262) (1:200) (Santa Cruz Biotechnology SC-67204)

Rat anti Ki-67 monoclonal antibody (SolA15), eFluor 570 (1:200) (eBioscience 41-5698-80)

Donkey anti-rabbit IgG Alexa Fluor 488 (Molecular Probes 1874771) (1:500)

Goat anti-Mouse IgG1 Alexa Fluor 488 (1:250) (Thermo Fisher Scientific A-21121)

Blocking buffer (5% goat serum in PBS with 0.3% Triton X)
