## Supplementary figures and images for "Microvascular preservation and cardiomyocyte hyperplasia underlie adaptive right ventricular development in congenital heart disease-associated pulmonary arterial hypertension"

### Supplemental Figure 1

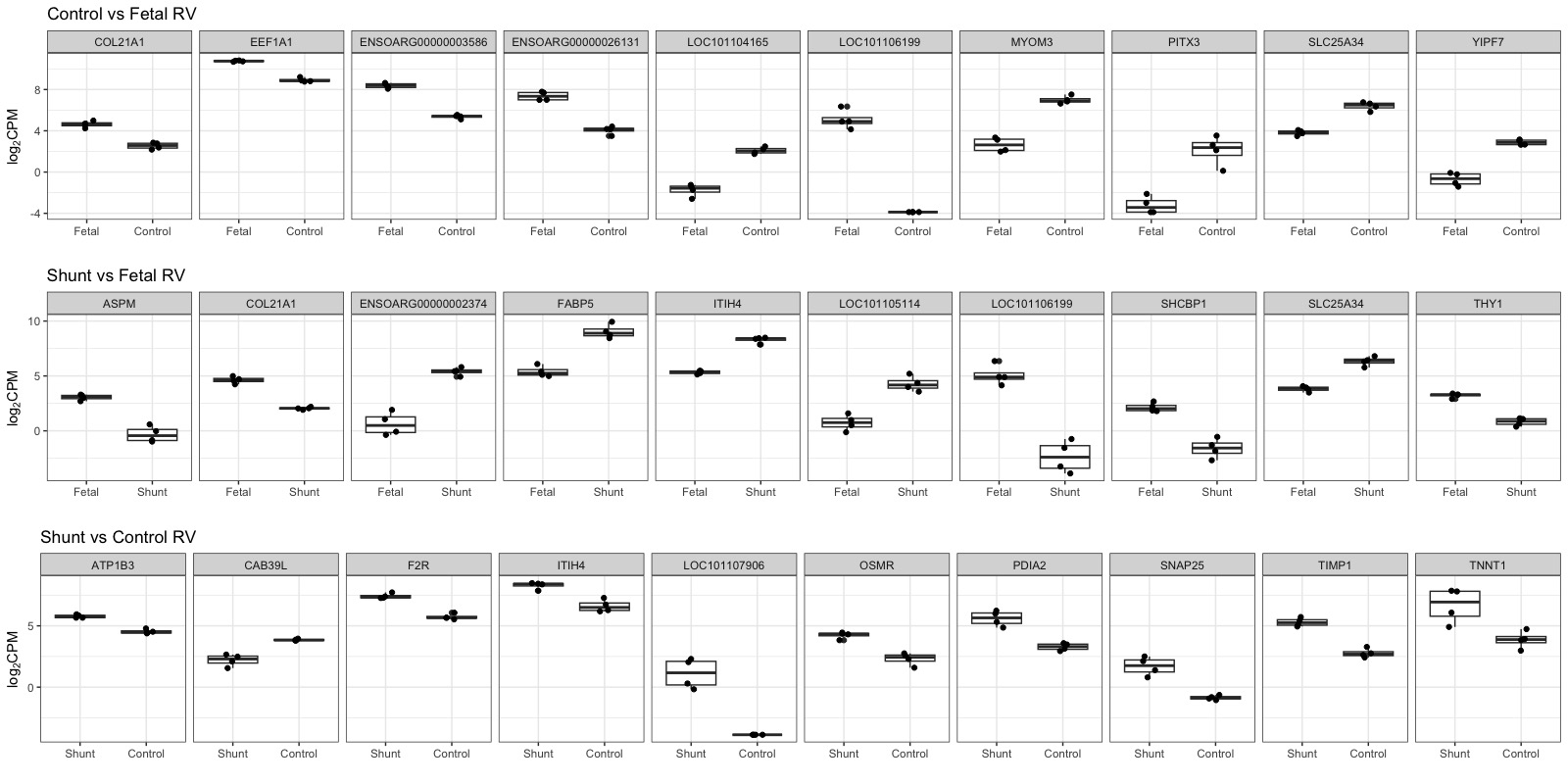
